# Pharmacological up-regulation of CALM1 restore endothelial cells in intracranial aneurysm pathogenesis

**DOI:** 10.64898/2026.08.04.742848

**Authors:** Danyi Zheng, Yikui Liu, Zhao Lanlan, Bing Leng, Qingfang Sun, Baofeng Wang, Xuanfeng Qin, Liuguan Bian, Yongtao Zheng

## Abstract

Subarachnoid hemorrhage (SAH) resulted from intracranial aneurysm (IA) rupture is an especially severe form of stroke. Endothelial dysfunction represents the initiating event of IA pathogenesis. Understanding the role of endothelial cells (ECs) underlying formation of IAs is helpful to seek for pharmaceutical treatment strategy. Based on single-cell RNA sequencing, proteomics, and metabolic analysis, we discovered a change in cell population in IA samples, majorly in ECs and macrophages (MPs). Abnormal ECs exhibit senescence and death in IA samples, which is absent in the control arterial samples. Cross-analysis of multi-omics revealed that CALM1, a calcium detector involved in mechanotransduction, is downregulated in the abnormal ECs. CALM1 knockdown leads to senescence and inhibits the proliferation and maturation of ECs under turbulent flow. Through high-throughput virtual screening, this work identified compound ZC04329651 as a potent CALM1 activator in maintaining the stability of endothelial cell junctions and attenuating cellular senescence. Thus, our findings showed compound ZC04329651 up-regulate the expression of CALM1 to restore ECs, which maybe a promising pharmacological treatment strategy for IAs.

## Introduction

Stroke ranks as the second most common cause of death globally, after ischemic heart disease. Subarachnoid hemorrhage (SAH) resulted from intracranial aneurysm (IA) rupture is an especially severe form of stroke, results in a mortality rate of approximately 30%-45%.^[1–3]^ Although surgical clipping and endovascular coiling are used for the prevention of aneurysmal rupture, potentially serious complications from the clipping and coiling of unruptured IAs are not neglected^[4–5]^. The management of unruptured IAs, especially very small aneurysms, remains a clinical dilemma, as preventive treatment carries inherent procedural risks, whereas conservative management may substantially impair quality of life due to the mental burden. These challenges emphasize the urgent need to develop effective non-invasive therapies for unruptured IAs.

Understanding the mechanisms underlying formation and rupture of IAs is helpful when seeking for effective non-invasive therapeutic strategies, for example pharmaceutical treatment. At arterial bifurcations, accelerated blood flow generates a unique hemodynamic environment characterized by high wall shear stress and a positive shear stress gradient along the flow direction. It activates intracellular signaling cascades via endothelial cell (EC)-mediated mechanotransudction, resulting in cell-cell junction disruption. The local vascular wall injury triggers inflammatory cell infiltration and inflammatory mediators release, followed by apoptosis and migration of smooth muscle cells (SMCs), remodeling and degradation of the extracellular matrix, ultimately leading to aneurysm formation and rupture.^[6–8]^ Endothelial dysfunction represents both the initiating event and a critical driver of IA pathogenesis. Consequently, exploring EC function during aneurysm progression is crucial for understanding IA’s pathogenic mechanism and provides insights for the development of non-invasive therapeutic strategies.

To elucidate the pathogenic mechanisms of aneurysm formation, we performed multi-omics studies to the human patient IA and control samples. Transcriptomic analysis revealed a change in cell population of IA samples, especially in ECs and macrophages. Cross-studies with proteomics analysis and metabolomic analysis revealed CALM1-induced senescence and death of ECs as a key factor in IA development. Using human iPSC-derived EC organoids, we showed that CALM1 knockdown leads to the senescence of ECs and inhibits the proliferation and maturation of the blood vessel organoids. These results pinpointed a CALM1-downregulation-mediated pathway in EC-triggered IA development, and opened a possibility of developing preventive and diagnostic strategies targeting CALM1 for cerebrovascular diseases. Further, through virtual screening and experimental validation, we identified a potent and specific agonist of CALM1, compound ZC04329651. Pharmacological upregulation of CALM1 by compound ZC04329651 alleviated cellular senescence of hiPSC-derived arterial ECs and promoted the formation of cell junctions. Overall, our study establishes CALM1 as a promising therapeutic target for IA and positions compound ZC04329651 as a lead compound for future chemical optimization in preclinical drug development.

## Methods

### Patients and samples

The protocol for collecting human tissue samples was approved by the Research and Ethical Committee of Ruijin Hospital, Shanghai Jiaotong University (Protocol number 2025-350), and complied with the Declaration of Helsinki. Written informed consent was obtained from all participants before enrolment. All patients were performed with digital subtraction angiography (DSA) before clipping. Patients were excluded if they had fusiform aneurysms, dissection aneurysms, and/or pseudoaneurysms. In total, five IA samples were dissected and obtained after the aneurysm neck was satisfied with surgical clipping (Table 1). For control purposes, five superficial temporal arteries (STAs) samples were obtained from those patients when the STA is inevitably injured during craniotomy.

### Single-cell RNA sequencing analysis

Single-cell RNA-seq experiment was performed by experimental personnel in the laboratory of NovelBio Co., Ltd. The tissues were removed and kept in MACS Tissue Storage Solution (Miltenyi Biotec) until processing. The tissue samples were processed as described below. Briefly, samples were first washed with 1640 culture solution, minced into small pieces (approximately 1mm3) on ice and enzymatically digested with Collagenase I 1 mg /ml (Worthington), Dispase 1 mg /ml (Worthington), DNase I 30 U/ml (Worthington) for 20 min at 37°C; Second round, digested with 3ml Trypsin for 20 min with agitation. After digestion, the pelleted cells were suspended in red blood cell lysis buffer (Miltenyi Biotec) to lyse red blood cells. After washing, the cell pellets were re-suspended in 1640 culture solution and then stained with AO/PI for viability assessment using Countstar Fluorescence Cell Analyzer.

BD Rhapsody system was used to capture transcriptomic information of the single cells. Single-cell capture was achieved by random distribution of a single-cell suspension across >200,000 microwells through a limited dilution approach. Beads with oligonucleotide barcodes were added to saturation so that a bead was paired with a cell in a microwell. Cell-lysis buffer was added so that poly-adenylated RNA molecules hybridized to the beads. Beads were collected into a single tube for reverse transcription. Upon cDNA synthesis, each cDNA molecule was tagged on the 5′ end (that is, the 3′ end of a mRNA transcript) with a unique molecular identifier (UMI) and cell label indicating its cell of origin. Whole transcriptome libraries were prepared using the BD Rhapsody single-cell whole-transcriptome amplification workflow. In brief, second strand cDNA was synthesized, followed by ligation of the WTA adaptor for universal amplification. Eighteen cycles of PCR were used to amplify the adaptor-ligated cDNA products. Sequencing libraries were prepared using random priming PCR of the whole-transcriptome amplification products to enrich the 3′ end of the transcripts linked with the cell label and UMI. Sequencing libraries were quantified using a High Sensitivity DNA chip (Agilent) on a Bioanalyzer 2200 and the Qubit High Sensitivity DNA assay (Thermo Fisher Scientific). All libraries were sequenced by DNBSEQ-T7 Sequencer (MGI, Shenzhen, China) on a 150 bp paired-end run.

scRNA-seq data analysis was performed by NovelBio Co.,Ltd. with NovelBrain Cloud Analysis Platform. We applied fastp with default parameter filtering the adaptor sequence and removed the low-quality reads to achieve the clean data. To quantify the gene expression of the single-cell data, we used STARsolo (version 2.7.10a) along with human genome GRCh38 (ensemble annotation version 104). Cells contained over 200 expressed genes and mitochondria UMI rate below 20% passed the cell quality filtering and mitochondria genes were removed in the expression table. All subsequent analysis were performed using R (version 4.5.1) package Seurat (version 5.0.3). The raw count matrices were normalized by natural-log transformation with a scale factor of 10000. Dimensionality reduction using principal component analysis (PCA) was performed, and clusters of nuclei were identified in principal component analysis (PCA) space by shared nearest-neighbor graph construction and modularity detection implemented by the FindNeighbors and FindClusters functions using a dataset dimension of 30 and resolution of 1.0. The dataset was embedded for visualization with UMAP. Marker genes and DEGs for each cluster were calculated using FindMarkers with default parameters based on normalized data and filtered by p-value < 0.05. To better understand the cell population, we integrated the dataset to the adult human cerebrovasculature snRNA-seq reference dataset (which can be downloaded through the UCSC Cell Browser: https://adult-brain-vasc.cells.ucsc.edu) using the Anchor-based CCA integration in Seurat. Only genes detected in both datasets were included in comparison. Our cells were defined using the label from the reference clusters where they were overlapped. Gene Ontology (GO) and KEGG pathway enrichment analysis were performed using enrichR.

### Proteomic and metabolomic sample preparation

Sample preparation was performed based on previously established proteomic methods. Briefly, samples were lysed with 100 μL of lysis buffer containing 1% sodium deoxycholate (SDC), 10 mM tris-(2-carboxyethyl)-phosphine (TCEP), 20 mM chloroacetamide (CAA), 0.1% RapiGest surfactant, and 1× protease inhibitor in 50 mM ammonium bicarbonate (ABC). Lysis was conducted at 95°C for 60 min, followed by ultrasonication for 10 cycles (30 s ON/OFF). For protein digestion, trypsin and Lys-C were added at enzyme-to-protein ratios of 1:50 (w/w) and 1:100 (w/w), respectively, and incubated at 37°C for 2 h with shaking at 900 rpm. The digestion was quenched by adding TFA to a final concentration of 2% and incubating at 37°C for 30 min, followed by centrifugation at 13,000 × g for 10 min. The supernatant was desalted, lyophilized to dryness, and either stored at −20°C or reconstituted in 0.1% formic acid (FA) for nano-LC-MS/MS analysis. Metabolic extracts were obtained using methanol-assisted protein precipitation. Cold 80% methanol (−20°C) was added to the samples, which were then stored at -20°C overnight. The mixture was centrifuged at 15,000 × g for 15 min at 4°C. The supernatant was collected and the centrifugation step was repeated until a clear supernatant was obtained. The supernatant was lyophilized to dryness and stored at −20 °C or redissolved in 10% acetonitrile (ACN) for nano-LC−MS/MS analysis.

### Proteomics and metabolic analysis

For proteomics analysis, all subsequent analysis were performed using R (version 4.5.1). Dimensionality reduction using PCA was performed, and the raw dataset was normalized using housekeeping genes. The DEPs were filtered by p-value < 0.05. Gene Ontology (GO) and KEGG pathway enrichment analysis were performed using enrichR.

For metabolic analysis, all subsequent analysis were performed using R (version 4.5.1). Dimensionality reduction using PCA was performed, and the differentially produced metabolites were filtered by p-value < 0.05.

### Immunofluorescence Staining of Human IA Tissues

Samples of IAs were taken from patients undergoing microsurgical clipping. Immediately after complete aneurysm clipping, the aneurysmal wall was harvested and fixed in 4% paraformaldehyde for 2-4 hours at 4°C. Tissues were cryoprotected in 30% sucrose, embedded in optimal cutting temperature (OCT) compound, and snap-frozen. Cryosections (20 μm thick) were cut using a cryostat and mounted on glass slides. Sections were rinsed three times with PBS, then blocked with 5% fetal bovine serum (FBS) in PBS for 60 minutes at room temperature. Primary antibodies against CD31 and CALM1 were applied and incubated overnight at 4°C. After washing with PBS, sections were incubated with fluorophore-conjugated secondary antibodies for 60 minutes at room temperature, followed by nuclear counterstaining with DAPI. Fluorescent images were acquired using a confocal laser scanning microscope.

### Cell culture and Transfection of shRNA

hiPSC-derived arterial ECs were maintained in DMEM supplemented with 10% FBS under standard conditions (37°C, 5% CO₂, 95% humidity). Cells were passaged every 48 hours upon reaching 80-90% confluence. To silence CALM1 expression, hiPSC-derived arterial ECs were transduced with a lentiviral vector (pLenti-U6-shRNA (CALM1)-CMV-BSR-WPRE; OBiO Biotech, Shanghai, China) according to the manufacturer’s protocol.

### hiPSC-Derived Vascular Organoids

Human-induced pluripotent stem cells (hiPSCs) were dissociated into single cells and gathered in a low-adhesion U-shaped 96-well plate (7,000-9,000 cells/well) in mTeSR1 medium containing 50uM Y-27632. The aggregates were cultured for 48 hours, then the mesoderm was induced by APEL2+6uM CHIR9902 for 2 days, and then the endothelium was induced by APEL2+50ng/ml VEGF+25ng/ml BMP4+10ng/ml bFGF for 2 days. After that, it was cultured in ECM basal medium containing 50ng/ml VEGF for 4 days to induce the maturation of ECs. On the 11th day, the aggregates were embedded in Matrigel and cultured in vascular maturation medium (ECM basal medium containing 50ng/ml VEGF). On the 13th day, the aggregates were dug into a low-adsorption 6-well plate and cultured in ECM basal medium containing 20ng/ml VEGF for 7 days. The quasi-organ was maintained for 20 days, and the medium was changed every 48 hours.

### EdU Assay

For proliferation assessment, hiPSC-derived arterial ECs were seeded on glass coverslips in 24-well plates (5× 10^4^ cells/well) and cultured overnight. Cells were pulsed with 10 μM EdU for 1 hour, then fixed with 4% paraformaldehyde and permeabilized with 0.5% Triton X-100. EdU incorporation was detected using the EdU-555 Cell Proliferation Kit (Beyotime, China) following the manufacturer’s instructions. Nuclei were counterstained with DAPI, and EdU-positive cells were visualized and quantified via fluorescence microscopy using ImageJ.

### Western Blot Analysis

Cells or tissue lysates were prepared in ice-cold RIPA buffer containing protease and phosphatase inhibitors. After centrifugation at 12,000 rpm for 10 minutes at 4°C, supernatants were collected, and protein concentration was determined. Equal amounts of protein (30 μg per lane) were resolved by SDS-PAGE and transferred onto PVDF membranes. Membranes were blocked with 5% non-fat milk in TBST for 1 hour at room temperature, then probed with primary antibodies overnight at 4°C. After washing, horseradish peroxidase (HRP)-conjugated secondary antibodies were applied for 60 minutes. Protein bands were visualized using enhanced chemiluminescence, and band intensities were quantified with ImageJ. All experiments were independently repeated at least three times.

### *In vitro* Hemodynamic Shear Stress Exposure

Laminar and turbulent flow conditions were generated using a Flexcell FX-6000T flow system (Flexcell International, USA). Slides (BF-3001C) were pre-coated with 50 μg/mL collagen I overnight. hiPSC-derived arterial ECs (5×10⁵ cells/slide) were seeded and allowed to adhere for 12 hours in complete medium. Laminar shear stress was applied at a constant 12 dyn/cm², while turbulent flow was simulated using an oscillatory waveform (0.5±6 dyn/cm², 1 Hz). After 6 hours of exposure, cells were either fixed for immunofluorescence or lysed for Western blot analysis.

### Molecular dynamics (MD) simulation and docking

For docking analysis, the CALM1 protein and the compound ZC04329651 molecule were converted into PDBQT formats using MGLTools. Subsequently, Autodock Vina was employed to dock the compound ZC04329651 molecule into the active site of CALM1, thereby generating the CALM1-compound ZC04329651 complex. MD simulations were performed using GROMACS 2022.2. The CALM1 protein was modeled with the Amber14SB force field, while the compound ZC04329651 ligand was parameterized using the GAFF2 force field. The complex was solvated with TIP3P water, maintaining a minimum distance of 1.2 nm between the protein and the box boundaries, and neutralized by adding Na^+^ and Cl^-^ ions. Energy minimization was conducted using the steepest descent method, which was followed by staged equilibration under NVT and NPT ensembles at 298 K for 200 ps each. Production MD simulations were then executed for 200 ns in the NPT ensemble. Finally, GROMACS and VMD/PyMOL were used to visualize the interaction patterns. The binding free energy was calculated via gmx_MMPBSA.

### High-throughput virtual screening (HTVS)

To identify potential CALM1 agonists, Schrödinger Maestro version 12.8 was used for structure-based HTVS. The X-ray crystal structure of human CALM1 (Protein Data Bank 7PSZ) was processed using the Protein Preparation Wizard. The MedChemExpress compound libraries (HY-L001P, HY-L901P, and HY-L0086V), containing 305,300 compounds, were prepared for virtual screening through energy optimization by the LigPrep module of Schrödinger Maestro. These optimized compounds were then subjected to a virtual screening workflow. First, all the compounds were screened in Glide HTVS mode, and the top compounds were subjected to Glide standard precision (SP) docking. The top SP results were subsequently docked in extra precision (EP). Based on the SP and EP scores, the top-ranked compounds from three libraries were selected: HY-L001P (ranked top 15%), HY-L901P (ranked top 10%), and HY-L0086V (ranked top 5%). According to documented bioactivity on endothelial cells, the 51 compounds were further screened as candidates for experimental validation. Following virtual screening, single-concentration surface plasmon resonance (SPR) assays were conducted to assess the binding affinity of these 51 compounds. For compound sitagliptin phenylalanine, multi-concentration SPR analysis was performed, and the dissociation constant (KD) for its binding to CALM1 was determined by globally fitting the sensorgrams to a 1:1 Langmuir binding model using the Biacore Insight Evaluation Software (Cytiva, Marlborough, MA, USA).

### Quantification and Statistical Analysis

The data of the stroke group and the transplanted groups are normally distributed. Unpaired t-test was used for comparison between two groups. Error bars in all figures represent means ± SEM. Differences were considered statistically significant at a p-value of < 0.05. For multi-group comparisons, one-way ANOVA followed by Tukey’s post hoc test was used. A p-value < 0.05 was considered statistically significant. All *in vitro* experiments were conducted in at least three independent biological replicates. All data were analyzed using GraphPad Prism 10.6.0.

## Results

### Multi-omics analysis reveals cellular senescence as a key factor in IA development

To study the pathological mechanisms of IA, we performed single-cell RNA sequencing, proteomics and metabolomics analysis to human patient samples. The single-cell RNA sequencing was performed using 5 IA samples (Aneurysm) and 1 STA sample (Ctrl). The cells were separated into 27 clusters, each with a set of distinct markers (Fig. 1a and Supplementary Fig. 1a). To better understand the cell population of human patient samples, we integrated the Aneurysm samples and the Ctrl sample to the adult human cerebrovasculature scRNA-seq reference dataset (WT) (Fig. 1b). The cells were separated into 45 clusters, each with a set of distinct markers (Supplementary Fig. 1b). Transferring the labels from the public dataset, the cells in the integrated dataset were categorized as 23 cell types (Fig. 1c and Supplementary Fig. 1c). For further analysis, we categorized the 23 cell types into 4 major cell types: ECs, Lymphocytes (LCs), Macrophages (MPs), and Perivascular cells (PCs). The distribution of four major types of the Ctrl group was more similar as the WT group, while the Aneurysm group showed a notably smaller proportion of ECs and larger proportion of MPs (Fig. 1c,d). Although the Aneurysm and Ctrl group both consist of 22 cell types except pericytes, the top marker genes of each cell type were similar to, but not the same as the WT group (Fig. 1e,f and Supplementary Fig. 1c). To minimize batch effects, we only used Aneurysm and Ctrl datasets in further analysis.

**Figure 1.**
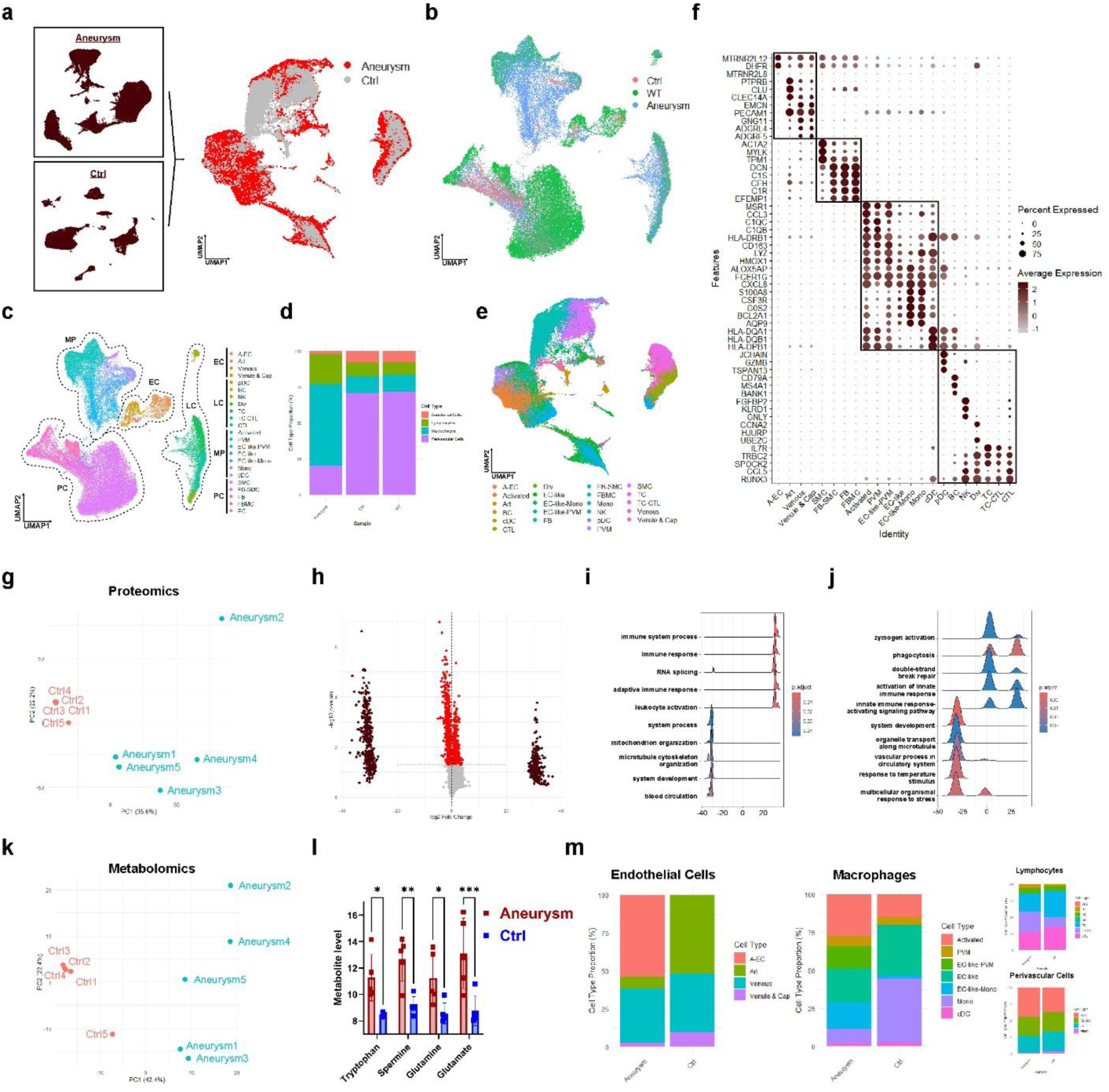
Multi-omics analysis reveals cellular senescence as a key factor in IA development. a, UMAP visualization of clustering of cells from human IA samples and Ctrl sample before (left) and after (right) Seurat integration. b, UMAP visualization of clustering of cells from human IA samples, Ctrl sample, and cells from adult human cerebrovasculature scRNA-seq reference dataset (WT) after Seurat integration. c, UMAP visualization of clustering of cells from human IA samples, Ctrl sample, and cells from the adult human cerebrovasculature scRNA-seq reference dataset (WT) after Seurat integration using labels transferred from the adult human cerebrovasculature scRNA-seq reference dataset. A-EC, abnormal endothelial cells; Art, arterial cells; pDC, plasmacytoid dendritic cells; BC, B cells; Nk, natural killer cells; Div, dividing lymphocytes; TC, T cells; CTl, cytotoxic T-lymphocytes; Activated, activated macrophages; PVm, perivascular macrophage; Mono, monocytes; cDC, conventional dendritic cells; SMC, smooth muscle cells; FB, fibroblasts; FBMC, fibromyocytes; PC, pericytes. d, Quantification of major cell type population of cells from human IA samples, Ctrl sample and WT, using labels transferred from the adult human cerebrovasculature scRNA-seq reference dataset. e, UMAP visualization of clustering of cells from human IA samples and Ctrl sample after Seurat integration using labels transferred from the adult human cerebrovasculature scRNA-seq reference dataset. f, Dotplot showing markers of each cell type in cells from human IA samples and Ctrl sample after Seurat integration. g, PCA plot for proteomics result of human IA samples and Ctrl samples. h, Volcano plot showing the differentially expressed proteins (DEPs, red) and “all-or-none” proteins (brown) of human IA samples and Ctrl samples. i, Ridgeplot showing Gene Ontology enrichment analysis result of “all-or-none” proteins of human IA samples and Ctrl samples. j, Ridgeplot showing Gene Ontology enrichment analysis result of DEPs and “all-or-none” proteins of human IA samples and Ctrl samples. k, PCA plot for the metabolomics result of human IA samples and Ctrl samples. l, Level of tryptophan, spermine, glutamine and glutamate from metabolomics result human IA samples and Ctrl samples. m, Quantification of cell population within each major cell type of cells from human IA samples and Ctrl sample.

The proteomics analysis was performed using 5 IA samples (Aneurysm) and 5 STA samples (Ctrl). PCA showed the Aneurysm and Ctrl samples are separated with a PC1 of 35.6%, with tight replicate clustering, indicating the consistent reproducibility of our proteomics approach (Fig. 1g). We identified 1185 differentially expressed proteins (DEPs) between the two groups, 637 of which were upregulated, and 548 of which were downregulated in the Aneurysm group. Gene Ontology (GO) enrichment analysis indicated that the upregulated DEPs were related to biosynthetic process, and the downregulated DEPs were related to the muscle function (Supplementary Fig. 1d). In addition, we also identified 794 “all-or-none” proteins, 359 of which were detected in all Aneurysm samples but in none of the Ctrl samples, and 435 of which were detected in all Ctrl samples but in none of the Aneurysm samples (Fig. 1h). GO enrichment analysis indicated that the “all” proteins in the Aneurysm group were related to immune response, and the “none” proteins were related to cellular activity and blood circulation (Fig. 1i). Taking together, GO enrichment analysis of the DEPs and “all-or-none” proteins showed that the Aneurysm group expressed more proteins related to immune response and senescence, and fewer proteins related to normal cellular function (Fig. 1j). The metabolomic analysis was performed 5 IA samples (Aneurysm) and 5 STA samples (Ctrl). PCA showed the Aneurysm and Ctrl samples are separated with a PC1 of 42.4%, with tight replicate clustering, indicating the consistent reproducibility of our metabolomic approach (Fig. 1k). We identified 4 metabolites, tryptophan, spermine, glutamine, and glutamate, which were related to cellular senescence, and were enriched in the Aneurysm group compared to the Ctrl group (Fig. 1l).

We then investigated the distribution of cell types in each major cell type. We found that cell distribution in the ECs and MPs was different between the Aneurysm group and Ctrl group, while that in the LCs and PCs was similar between the two groups. In the ECs, the Aneurysm group contains a majority of abnormal ECs (A-ECs), while the Ctrl group contains mainly normal blood vessel cells including artery (Art), venous, and venule & capillary cells (Fig. 1m). Together, the proteomics and metabolomics analysis indicated that cellular senescence is a key factor during aneurysm development, and single-cell RNA sequencing analysis indicated that cellular population change was mainly in ECs and MPs. This finding is line with the conventional understanding that senescence and death of ECs represent the initial trigger of aneurysm.

### Immune cells display cell fate shift, impaired cellular functions, and dysregulated immune responses in IAs

As discussed in the previous section, the cell population of MPs was different between the Aneurysm group and Ctrl group. We identified a few clusters of cells which exhibited a transition in identity from a common cell type to a hybrid state, which we termed as EC-like, EC-like-Mono, and EC-like-PVM cells (Fig. 1m and Fig. 2a). The EC-like cells were clustered close to ECs, although remain in MPs on the UMAP. Compared to ECs, EC-like cells express a higher level of genes related to immune response, and a lower level of genes related to angiogenesis (Fig. 1c and Supplementary Fig. 2a). The EC-like-PVM cells express a higher level of genes related to MHC protein complex, and a lower level of genes related to extracellular matrix (ECM) compared to EC-like cells. They also express a higher level of immune response-associated genes and a lower level of cell junction-related genes compared to PVM cells (Supplementary Fig. 2b). The EC-like-Mono cells express a higher level of genes related to immune response and a lower level of genes related to ECM. They also express a higher level of genes related to immune activation, and a lower level of genes related to metabolic processes compared to monocytes (Mono) (Supplementary Fig. 2c). These intermediate cells mainly appeared in the Aneurysm group but not in the Ctrl group, indicating a cell fate change in the IA samples (Supplementary Fig. 2d). We investigated the differentially expressed genes (DEGs) between the Aneurysm and Ctrl group within each cell type and found that DEGs derived from each of the cell type frequently exhibited similar expression trends across other cell types (Fig. 2b). Similarly, GO enrichment analysis revealed that upregulated genes in each cell type were predominantly associated with biosynthetic and metabolic processes, whereas downregulated genes were enriched for terms related to immune responses and cell fate specification (Fig. 2c).

**Figure 2.**
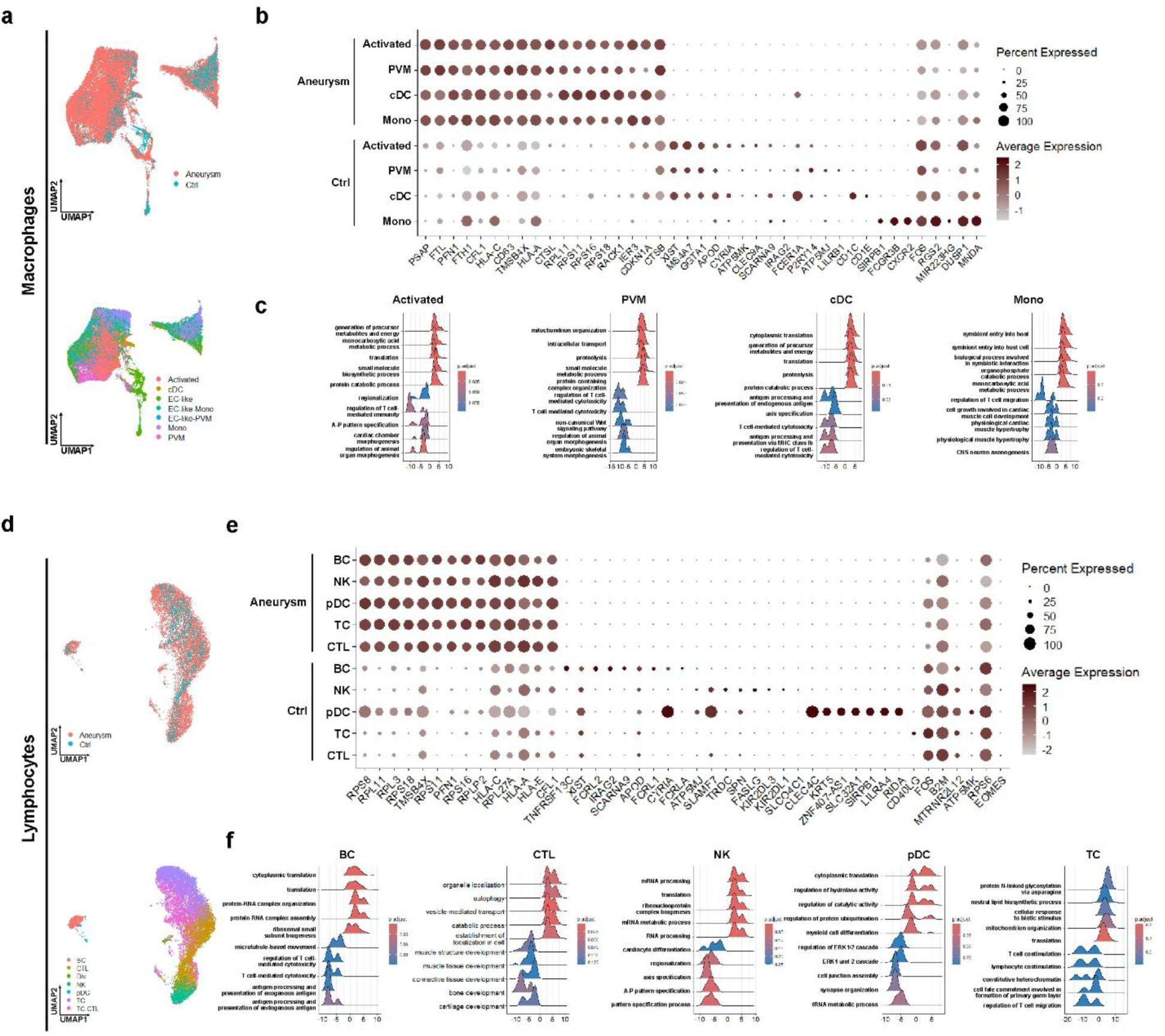
Immune cells display cell fate shift, impaired cellular functions, and dysregulated immune responses in IAs. a, UMAP visualization of clustering of macrophages from human IA samples and Ctrl sample after Seurat integration using labels transferred from the adult human cerebrovasculature scRNA-seq reference dataset. b, Dotplot showing differentially expressed genes (DEGs) between human IA samples and Ctrl sample within each cell type in macrophages. c, Ridgeplots showing Gene Ontology enrichment analysis of DEGs between human IA samples and Ctrl sample within each cell type in macrophages. d, UMAP visualization of clustering of lymphocytes from human IA samples and Ctrl sample after Seurat integration using labels transferred from the adult human cerebrovasculature scRNA-seq reference dataset. e, Dotplot showing differentially expressed genes (DEGs) between human IA samples and Ctrl sample within each cell type in lymphocytes. f, Ridgeplots showing Gene Ontology enrichment analysis of DEGs between human IA samples and Ctrl sample within each cell type in lymphocytes.

The cell population of LCs was similar between the Aneurysm and Ctrl group. We also identified a cluster of hybrid state, which we termed as TC-CTL cells (Fig. 1m and Fig. 2d). These cells express a higher level of genes related to cell migration, and a lower level of genes related to T-cell differentiation compared to T-cells (TC). They also express a higher level of genes related to blood vessel identity and a lower level of genes related to MHC protein complex compared to cytotoxic T lymphocytes (CTL) (Supplementary Fig. 2e). TC-CTL cells mainly appeared in the Aneurysm group but not in the Ctrl group, indicating a cell fate change in the IA samples (Supplementary Fig. 2f). We investigated the DEGs between the Aneurysm and Ctrl group within each cell type. Similar as MPs, the DEGs derived from each of the cell types frequently exhibited similar expression trends across other cell types (Fig. 2e). GO enrichment analysis also revealed that upregulated genes in each cell type were predominantly associated with biosynthetic and metabolic processes, whereas downregulated genes were enriched for terms related to immune responses and cell fate specification (Fig. 2f). Together, the molecular profile of MPs and LCs indicated that the immune cells in the IA samples underwent shifts in cell fate, impaired cellular functions, and dysregulated immune responses, which is consistent to known findings of increased immune response in IA patient brains.

### Perivascular cells undergo pathological changes that disrupt vascular homeostasis in IAs

Perivascular cells are essential in vascular homeostasis and angiogenesis, including majorly SMCs, fibroblasts (FBs), as well as fibromyocytes (FBMCs). We identified one intermediate-state cluster which we termed as FB-SMC. Compared to SMCs, FB-SMC cells express a higher level of genes related to immune response and a lower level of genes related to chromosome segregation. They also express a higher level of genes associated with metabolic process, and a lower level of genes related to organ development (Supplementary Fig. 3a). Unlike immune cells, the cell population of PCs in the Aneurysm and Ctrl groups were similar, including the intermediate-state cell proportion (Fig. 1m, Fig. 3a, and Supplementary Fig. 3b). We investigated the DEGs between the Aneurysm and Ctrl group within each cell type. Similar as the immune cells, the top DEGs derived from each of the cell types frequently exhibited similar expression trends across other cell types (Fig. 3c). GO enrichment analysis of DEGs in SMCs revealed that the upregulated genes were enriched for proliferation, whereas the downregulated genes were related to adrenergic receptor signaling pathway, which is essential for vascular muscle contraction, indicating impaired muscle function. For the rest of the cell types, GO enrichment analysis revealed that upregulated genes in these cell types were predominantly associated with immune response, whereas downregulated genes were enriched for terms related to cell fate specification (Fig. 3d). Together, the molecular profile of perivascular cells indicated that the PCs underwent pathological changes in IAs, disrupting the vascular homeostasis.

**Figure 3.**
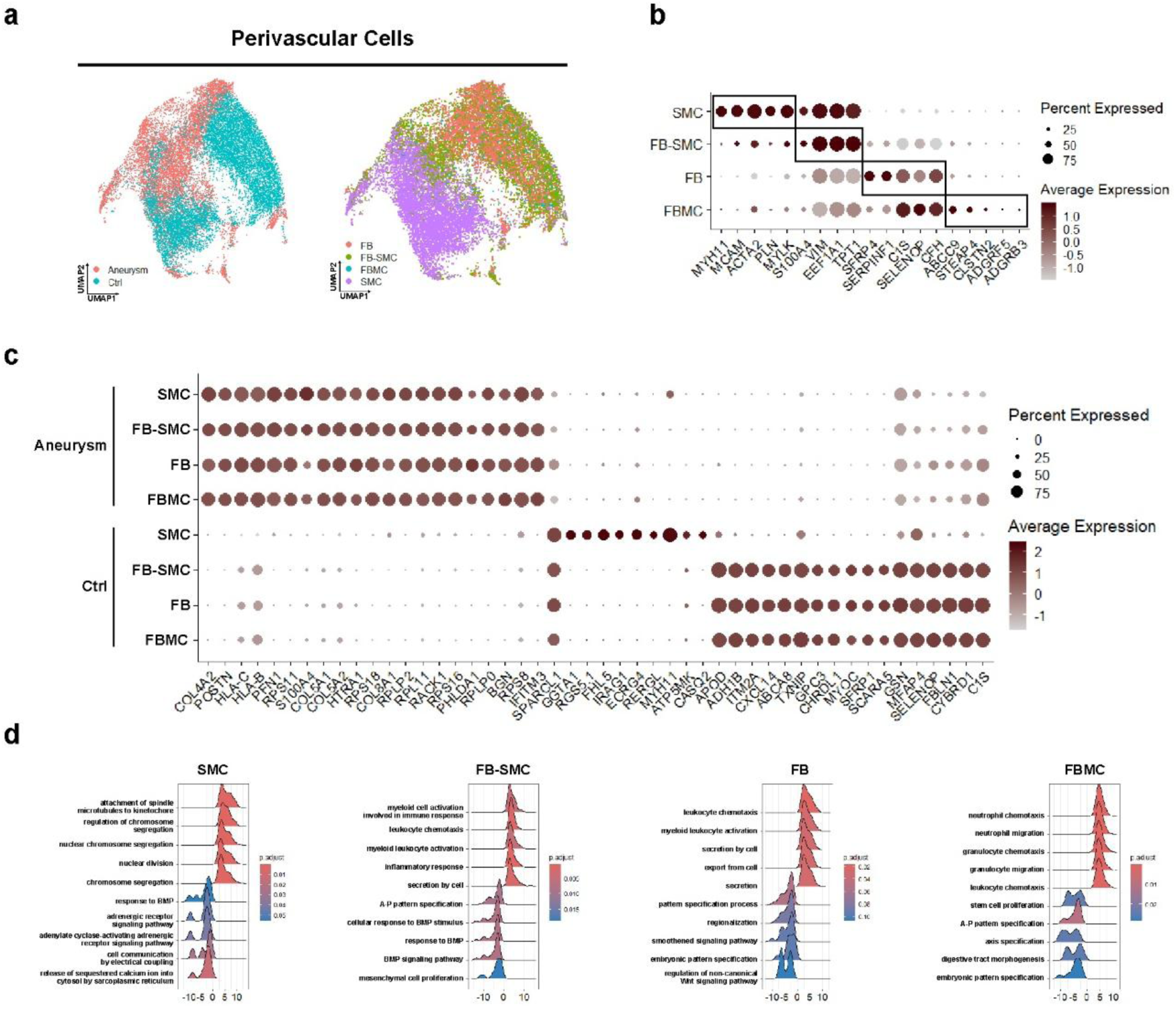
Perivascular cells undergo pathological changes that disrupt vascular homeostasis in IAs. a, UMAP visualization of clustering of perivascular cells from human IA samples and Ctrl sample after Seurat integration using labels transferred from the adult human cerebrovasculature scRNA-seq reference dataset. b, Dotplot showing marker genes of each cell type in perivascular cells from human IA samples and Ctrl sample after integration. c, Dotplot showing differentially expressed genes (DEGs) between human IA samples and Ctrl sample within each cell type in perivascular cells. d, Ridgeplots showing Gene Ontology enrichment analysis of DEGs between human IA samples and Ctrl sample within each cell type in perivascular cells.

### Endothelial cells exhibit senescence and cell death in IAs

As discussed in the previous section, the proportion of ECs was notably lower in the Aneurysm compared to the Ctrl group. Within the ECs, we identified a cluster of cells distinct from the common cell type which we termed as A-EC. A-ECs were detected exclusively in Aneurysm group and were absent from Ctrl group (Fig. 1d,m and Fig. 4a). These cells expressed a significantly higher level of mitochondrial genes, indicating the cells were experiencing cell stress and underwent apoptosis (Fig. 4b). Other than mitochondrial genes, A-ECs also expressed a lower level of anti-apoptosis genes such as NPM1, YBX1, etc (Fig. 4c). GO enrichment analysis revealed that the upregulated DEGs in A-ECs were mainly related to biosynthetic processes, whereas the downregulated DEGs were associated with vessel development and cell-cell adhesion (Fig. 4d). Together, these results indicated that A-ECs in the Aneurysm group experience cellular stress and exhibit features of cell death.

**Figure 4.**
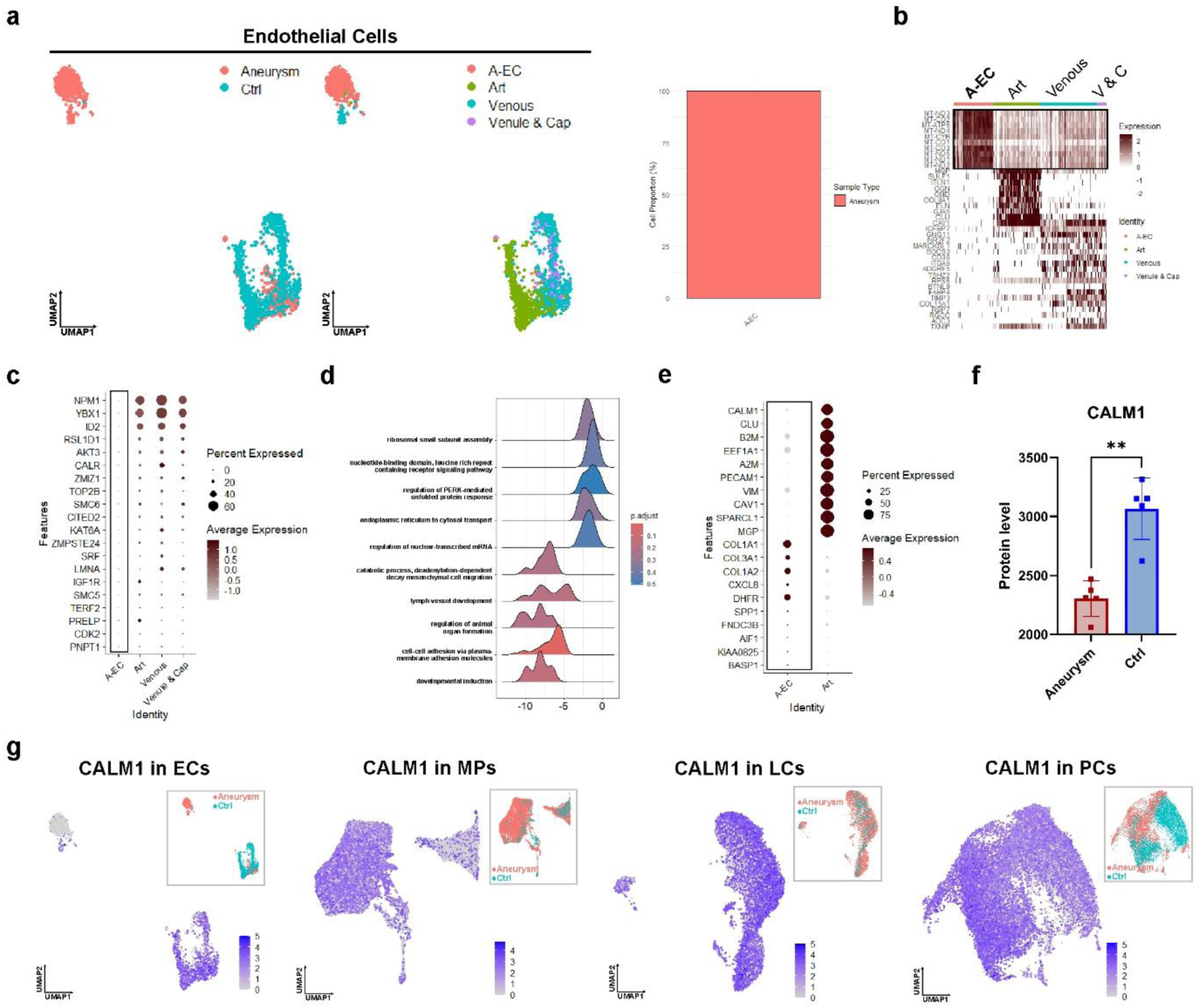
Endothelial cells exhibit senescence and cell death in IAs. a, UMAP visualization of clustering of endothelial cells from human IA samples and Ctrl sample after Seurat integration using labels transferred from the adult human cerebrovasculature scRNA-seq reference dataset (left). Quantification of distribution of A-EC in human IA samples and Ctrl sample (right). b, Heatmap showing marker genes of each cell type in endothelial cells from human IA samples and Ctrl sample after integration. c, Dotplot showing differentially expressed genes (DEGs) between A-EC and other endothelial cells. d, Ridgeplots showing Gene Ontology enrichment analysis of DEGs between A-EC and other endothelial cells. e, Dotplot showing differentially expressed genes (DEGs) between A-EC and normal arterial cells. f, Expression level of CALM1 in proteomics results of human IA samples and Ctrl samples. g, Feature plot showing expression level of CALM1 in each major cell type of cells from human IA samples and Ctrl sample after Seurat integration.

Interestingly, through integrated cross-comparison of transcriptomic and proteomic datasets, we found that the marked downregulation of CALM1 in these apoptotic cells is cell-type specific, occurring exclusively in endothelial cells (Fig. 4e-g). Notably, CALM1 is a key membrane protein mediating Ca²⁺ signaling, which is directly linked to mechanotransduction in endothelial cells. We therefore hypothesize that loss of CALM1 disrupts endothelial mechanosensing, thereby predisposing endothelial cells to senescence and apoptosis under abnormal blood flow.

### CALM1 knockdown promotes senescence, inhibits proliferation and blood vessel maturation in brain vessel organoids

To assess whether disturbed hemodynamics correlates with reduced CALM1 expression in human vascular pathology, we first analyzed CALM1 protein levels in clinical specimens from the walls of human Aneurysm samples and Ctrl samples. Quantification of CALM1 immunostaining fluorescence density showed that the CALM1 level in the Aneurysm samples was significantly lower than that of the Ctrl samples (Fig. 5a,d). The downregulation of CALM1 in Aneurysm samples was further supported by quantification of Western Blot (Fig. 5b).

**Figure 5.**
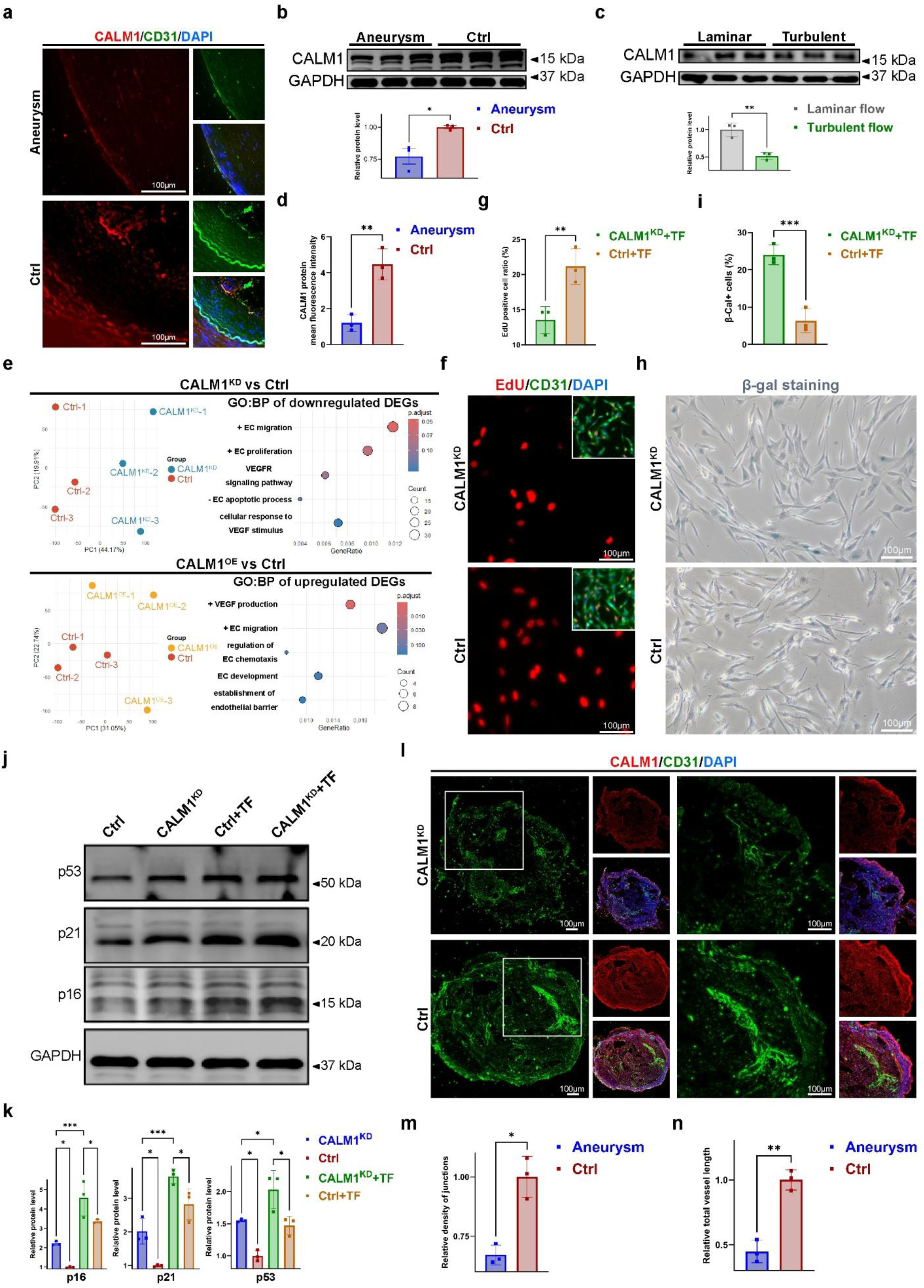
CALM1 protects hiPSC-derived arterial ECs against turbulent flow induced dysfunction. a, Immunofluorescence staining of CALM1 and endothelial marker CD31 in STA and IA tissue sections. b, Western blot analysis of CALM1 protein expression in human STA and IA tissues. c, Western blot analysis of CALM1 protein levels in hiPSC-derived arterial ECs exposed to laminar or turbulent flow for 6 h. Quantification shows significant downregulation of CALM1 under turbulent flow. d, Quantification of immunofluorescence density in Figure 5a. e, Bulk RNA-sequencing analysis results showing CALM1 has an important role in EC functions and prevents ECs from apoptosis. f, EdU incorporation assay assessing proliferation of control and CALM1^KD^ hiPSC-derived arterial ECs under turbulent flow. g, Quantification of EdU positive cells in Figure 5f. h, SA-β-galactosidase staining assessing senescence of CALM1^KD^ and control hiPSC-derived arterial ECs under turbulent flow. I, Quantification of SA-β-galactosidase cells in Figure 5h. j, Western blot analysis of senescence markers (p53, p21, p16) in CALM1^KD^ and control hiPSC-derived arterial ECs under turbulent flow. k, Quantification of Western Blot in Figure 5j. l, Immunofluorescence staining of CALM1 and endothelial marker CD31 in vascular organoids formed by CALM1^KD^ and control hiPSC-derived arterial ECs. m,n, Quantification of immunostaining in Figure 5l.

To determine whether turbulent flow directly leads to CALM1 downregulation, we treated *in vitro* cultured hiPSC-derived arterial ECs with defined laminar or turbulent flow conditions. Quantification of Western Blot showed that CALM1 protein level was significantly lower in hiPSC-derived arterial ECs exposed to turbulent flow compared to those exposed to laminar flow (Fig. 5c), suggesting that disturbed flow may cause EC injury. To investigate the role of CALM1 in ECs, we knocked down and overexpressed CALM1 in hiPSC-derived arterial ECs, respectively, and performed bulk RNA-sequencing analysis. Principal component analysis (PCA) showed that the CALM1^KD^ ECs and control (Ctrl) ECs are separated with a PC1 of 44% variance and the CALM1^OE^ ECs and Ctrl ECs are separated with a PC1 of 31% variance, with tight replicate clustering, indicating the consistent reproducibility of our RNA-sequencing approach. GO:Biological Process (GO:BP) enrichment analysis indicated that the downregulated genes in CALM1^KD^ ECs are related to positive regulation of EC migration, proliferation, VEGFR signaling pathway, and negative regulation of EC apoptosis. Similarly, GO:BP enrichment analysis indicated that upregulated genes in CALM1^OE^ ECs are related to positive regulation of VEGF production, EC migration, chemotaxis, development and establishment of endothelial barrier (Fig. 5e). Thus, we concluded that CALM1 in ECs has an important role in EC functions and prevents ECs from apoptosis.

To validate our findings from RNA-sequencing analysis, we assessed ECs’ functions and senescence in hiPSC-derived arterial ECs under turbulent flow. Quantification of EdU immunostaining showed that percentage of EdU^+^ cells was significantly lower in CALM1^KD^ ECs. This indicated that proliferation is significantly reduced in CALM1^KD^ ECs compared to Ctrl hiPSC-derived arterial ECs (Fig. 5f,g). Quantification of SA-β-galactosidase staining percentage of β-gal^+^ cells was significantly higher in CALM1^KD^ ECs. This indicated that senescence was significantly increased in CALM1^KD^ ECs (Fig. 5h,i). Quantification of Western Blot indicates that CALM1^KD^ hiPSC-derived arterial ECs expressed a higher level of p53, p21, and p16 compared to the Ctrl group, and exposure to turbulent flow further increased expression level of these proteins, indicating CALM1 knockdown accelerated cellular senescence of hiPSC-derived arterial ECs (Fig. 5j,k).

To better mimic the situation in human aneurysm, we induced 3D-cultured human vascular organoids *in vitro*. Quantification of immunostaining showed that density of cell-cell junctions and total brain vessel length were significantly reduced in CALM1^KD^ organoids (Fig. 5l-n). Collectively, these results indicated that turbulent flow downregulates CALM1, and further knockdown of CALM1 exacerbates endothelial dysfunction, suppressing proliferation and migration while promoting senescence and impairing vascular morphogenesis. These results suggested that the downregulation of CALM1 is unlikely to be a consequence of endothelial senescence. Instead, reduced CALM1 expression renders endothelial cells more susceptible to senescence and even apoptosis under hemodynamic stress. Notably, restoration of CALM1 expression effectively protects endothelial cells from flow-induced senescence and apoptosis. This finding identifies CALM1 as a potential therapeutic target for the prevention and treatment of IAs.

### Virtual screening and experimental validation identified CALM1 agonists

Given the important role of CALM1 in promoting endothelial cell repair of IAs, we sought to identify small molecule CALM1 agonists. Through HTVS of 305,300 small molecule compounds from three MedChemExpress libraries, we identified chemical compounds capable of binding to CALM1 (Fig. 6a). Based on the SP and XP docking results, 51 candidate compounds relevant to endothelial functions were selected for further affinity screening via SPR. SPR screening identified 17 compounds with stronger binding affinities (RU>20) (Fig. 6b). Subsequently, we examined the effects of these 17 compounds on CALM1 expression in ECs. Quantification of Western Blot showed that only compound ZC04329651 significantly upregulated CALM1 expression, indicating that compound ZC04329651 is a potential up-regulator of CALM1 (data not shown).

**Figure 6.**
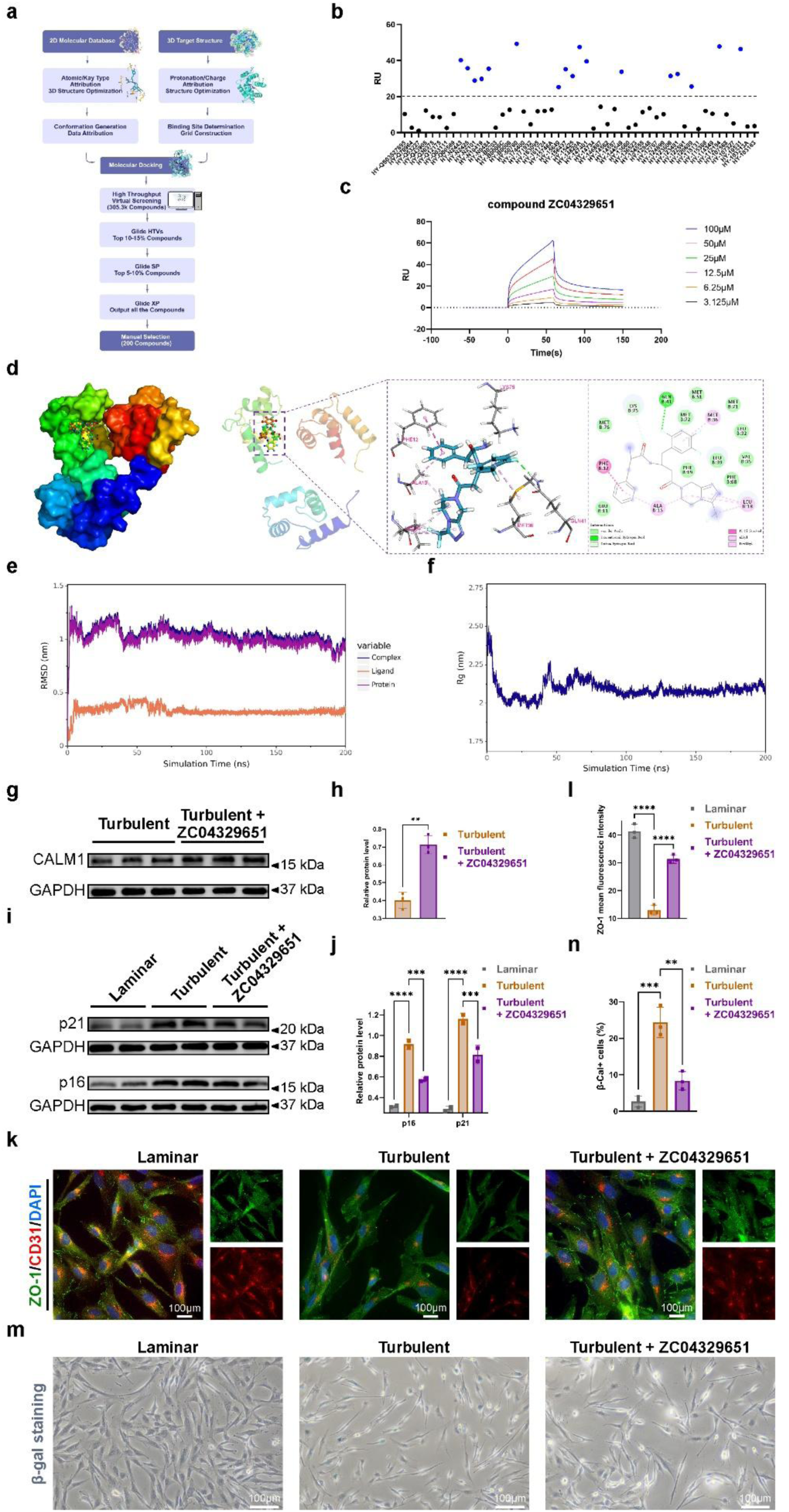
Virtual screening and experimental validation identified CALM1 agonists. a, Workflow of virtual screening and validation of CALM1 agonists. b, Single-concentration SPR screening against CALM1 protein to identify candidate compounds with strong binding affinities. c, Binding affinity of compound ZC04329651 to CALM1 determined by SPR. d, Molecular docking of compound ZC04329651 within the active pocket of CALM1. e, RMSD of the CALM1-compound ZC04329651 complex. f, Rg of the CALM1-compound ZC04329651 complex. g,h, Quantification of western blot analysis shows that compound ZC04329651 up-regulates the level of CALM1 in HUVECs under turbulent flow. i,j, Western blot analysis shows compound ZC04329651 inhibited cellular senescence of HUVECs under turbulent flow. k,l, Immunofluorescence staining indicates the intensity of ZO-1 was restored by the treatments of compound ZC04329651 under turbulent flow. m, n, Quantification of SA-β-galactosidase staining showed that compound ZC0432951 decrease the percentage of β-gal^+^ cells under turbulent flow.

SPR assay further confirmed that compound ZC04329651 directly binds to CALM1 protein with a KD value of 60.2 µM (Fig. 6c). Additionally, molecular docking also revealed a stable interaction between compound ZC04329651 and CALM1, with a binding affinity of -7.4 kcal/mol (Fig. 6d). GLN41 of CALM1 formed the hydrogen bond connections with compound ZC04329651. The dynamic changes in the compound ZC04329651-CALM1 interaction were assessed by MD simulations, with stable root-mean-square deviation (RMSD) and radius of gyration (Rg) during the 75-200 ns range (Fig. 6e,f). The binding free energy was -187.345±5.917 kJ/mol. These results suggest that compound ZC04329651 stably binds to CALM1.

Subsequently, to evaluate the effect of compound ZC04329651 on endothelial cell activity, we subjected cultured ECs to different flow conditions *in vitro* and applied with ZC0432951. Quantification of Western Blot showed ZC0432951 increased protein level of CALM1 in ECs under turbulent flow (Fig. 6g,h). Quantification of Western Blot showed that the expression level of p21 and p16 was increased in ECs under turbulent flow compared to those under laminar flow, and ZC0432951 alleviated the increase, indicating ZC04329651 inhibited cellular senescence of hiPSC-derived arterial ECs (Fig. 6i,j). Quantification of immunostaining showed that ZO-1 was decreased in ECs under turbulent flow compared to those under laminar flow, and ZC0432951 alleviated the decrease, indicating that disrupted cell-cell adhesion of ECs by turbulent flow was partially restored with ZC0432951 treatment (Fig. 6k,l). Quantification of SA-β-galactosidase staining showed that percentage of β-gal+ cells was significantly higher in ECs under turbulent flow compared to those under laminar flow, and ZC0432951 alleviated the increase, indicating ZC0432951 inhibited cellular senescence of ECs by turbulent flow (Fig. 6m,n).

Together, these results suggest that compound ZC04329651 has a relatively strong binding affinity with CALM1 and contributes to maintaining the stability of endothelial cell junctions and attenuating cellular senescence.

## Discussion

To systematically reveal the cell heterogeneity and risk factors underlying the development of IA, we conducted single-cell RNA sequencing, proteomics, and metabolic analysis of human IA samples. Our results indicated the critical roles of ECs and macrophages in the development of IA, and more specifically, a subpopulation of abnormal ECs featuring senescence and cell death. Integration of scRNA-seq and proteomics analysis demonstrated that CALM1 is specifically downregulated protein in IA ECs. Knockdown of CALM1 in brain vessel organoids results in suppressed proliferation and migration, as well as increased senescence and impaired vascular morphogenesis, further proving the important role of CALM1 in IA EC pathogenesis. Furthermore, we found that ZC04329651 increased CALM1 expression in ECs, and *in vitro* experiments validated ZC04329651 treatment promoted proliferation and cell-cell adhesion of ECs and inhibited EC senescence under turbulent flow.

IAs are a common cerebrovascular disorder affecting approximately 3–9% of the general population^[13–14]^. Although microsurgical clipping and endovascular intervention have substantially reduced the risk of aneurysm rupture, these invasive procedures are associated with procedure-related complications, aneurysm recurrence, and long-term neurological deficits. Currently, there are no pharmacological therapies to reduce the risk of aneurysm rupture. The lack of effective medical treatment represents a critical unmet clinical need in the management of IAs. A thorough understanding of the pathogenesis of IAs is conducive to the development of medical therapies for their treatment.

In quiescent and structurally stable blood vessels, ECs typically form a cobblestone-like monolayer lining the luminal surface of the vasculature. They form a selectively permeable barrier via opening and closing of intercellular junctions. Beyond serving as a barrier, vascular ECs also regulate vascular remodeling by secreting pro-and anti-angiogenic factors, and vascular SMC contraction via secreting vasodilators and vasoconstrictors^[15–17]^. In addition, they mediate inflammatory responses through the release of chemokines and cytokines. Aberrant hemodynamic forces modulate gene expression, resulting in morphological change and dysfunction of ECs.^[18–19]^ Disturbed hemodynamics cause cellular shrinkage and detachment of ECs, led to the disruption of intercellular junctions and increased cell-cell distance, resulting in increased blood vessel permeability.^[20–21]^ Moreover, impaired ECs exhibit reduced nitric oxide synthase activity, compromising vascular tone and vasomotor function, and secrete inflammatory mediators which promote inflammatory response.^[22–27]^ Thus, the dysfunction and apoptosis of ECs represent the earliest driving forces in IA formation. In line with the current understanding, our analysis also indicated that ECs in IA samples experience cellular stress, exhibit senescence and cell death. This finding further supports the current understanding and leads us to look into detailed mechanisms underlying EC pathogenesis.

Single-cell RNA sequencing provides important insights for identifying key cell subpopulations that drive the pathogenesis and progression of IAs. By further integrating proteomics, we analyzed the pathology-related proteins specifically secreted by these cells. Interestingly, we found that CALM1 was significantly downregulated in vascular ECs of IAs, while no notable differences were observed in other cell types. This suggests that down-regulation of CALM1 in ECs may contribute to the development and progression of IAs.

As a key calcium-binding signaling protein, CALM1 converts intracellular Ca^2+^ level change into signals, which further regulates diverse cellular activities including proliferation, apoptosis, inflammation, and vascular smooth muscle contraction. In vascular ECs, CALM1 promotes endothelial nitric oxide synthase phosphorylation and accounts for approximately 80% of nitric oxide (NO) production^[28–30]^. Currently, there is lack of research investigating the relationship between CALM1 and the mechanisms underlying IA formation. Our studies proved that downregulation of CALM1 induces endothelial senescence and apoptosis, suggesting that CALM1 deficiency may represent a key factor in endothelial dysfunction in IAs. Interestingly, the demonstrated protective role of CALM1 is consistent in other vascular diseases ^[31]^. In diabetic models, local CALM1 gene delivery improves post-ischemic blood perfusion and angiogenesis, further supporting CALM1 as a critical regulator of vascular endothelial homeostasis.

Using a human iPSC-derived vascular organoid model, we further demonstrate that CALM1 deficiency disrupts three-dimensional network formation, resulting in fewer junctions and shorter vessel segments—features reminiscent of early aneurysm pathogenesis. This organoid system offers advantages over conventional 2D cultures by better recapitulating endothelial self-organization and lumen formation, although it lacks physiological flow, mural cells, and immune components present *in vivo*. Notably, due to technical and resource limitations, our study lacks *in vivo* validation. Nevertheless, our vascular organoid model clearly demonstrates that CALM1 is essential for endothelial network integrity under hemodynamic stress. Together, these results suggest that CALM1 down-regulation in regions exposed to disturbed flow may contribute to endothelial vulnerability in IAs, highlighting its potential as a therapeutic target. Next, we performed drug screening and identified compound ZC04329651 which binds to CALM1 and upregulates CALM1 expression in ECs. ZC04329651 application on ECs was shown to promote EC proliferation and cell-cell adhesion, as well as inhibit cellular senescence, preserving vascular wall integrity.

Our findings highlight the translational potential of targeting CALM1 for the treatment of IAs. In contrast to current pharmacological approaches such as statins, matrix metalloproteinase inhibitors, and anti-inflammatory agents, ZC04329651 acts through a distinct molecular target and mechanism, avoiding ubiquitous side effects. As a small-molecule activator, ZC04329651 meets drug development needs, including convenient administration, flexible dose optimization, and potentially lower manufacturing costs compared with gene therapy–based approaches. More importantly, because endothelial dysfunction represents one of the earliest pathogenic events in IA development, targeting CALM1 may not only slow aneurysm progression but also provide an opportunity for preventive intervention during early stages, reducing the need for invasive treatment.

There are some limitations in this study. There are two types of IA, a thick-walled phenotype driven by high wall shear stress and dominated by inflammatory cell infiltration, and a thin-walled phenotype driven by low wall shear stress and characterized primarily by endothelial injury. As these two subtypes exhibit distinct pathogenic mechanisms, future studies should incorporate aneurysms of different types and anatomical locations to more comprehensively elucidate the mechanisms of aneurysm formation. Additionally, *in vivo* studies are needed to investigate the effective concentration range within aneurysmal lesions as well as long-term safety and efficacy.

To conclude, we are the first to integrate single-cell transcriptomics, proteomics, and metabolomics analyses of human IA samples. This multimodal approach enabled the novel discovery that reduced CALM1 expression in vascular endothelial cells is closely associated with aneurysm initiation and progression. We further demonstrated that CALM1 knockdown under disturbed-flow conditions impairs endothelial proliferation and migration while promoting apoptosis *in vivo*. We also pioneered the application of an organoid model to IA research, establishing an integrated experimental workflow that combines multi-omics profiling, hemodynamic simulation, and organoid technology. This strategy provides a new conceptual framework for dissecting the pathophysiological mechanisms of IAs. More importantly, our study identified CALM1 as a novel insight for IA treatments. We also identified compound ZC0432951 as a potential small molecule drug for IA prevention, which upregulates CALM1 expression and improves cell-cell adhesion of ECs and inhibited EC senescence.

## Acknowledgements

1. D. Z. and Y.Z. conceptualized and designed the experiments; Y.L., B.L., Q.S. and B.W. performed data collection; D.Z., Y.L., X.Q. and L.Z. performed data analysis and interpretation; D.Z., L.B. and Y.Z. drafted the article.

## References

1. Wessels, L., Wolf, S., Adage, T., Breitenbach, J., Thomé, C., Kerschbaumer, J., Bendszus, M., Gmeiner, M., Gruber, A., Mielke, D., et al. Localized Nicardipine Release Implants for Prevention of Vasospasm After Aneurysmal Subarachnoid Hemorrhage: A Randomized Clinical Trial. JAMA Neurol 81, 1060–1065 (2024) .

2. Zheng, Y., Xu, F., Ren, J., Xu, Q., Liu, Y., Tian, Y., Leng, B. Assessment of intracranial aneurysm rupture based on morphology parameters and anatomical locations. J Neurointerv Surg 8, 1240–1246 (2016) .

3. Engel, A., Song, L., Rauschenbach, L., Gümüs, M., Santos, A.N., Dinger, T.F., Darkwah Oppong, M., Li, Y., Gembruch, O., Ahmadipour, Y., et al. Impact of Carotid Siphon Calcification on the Course and Outcome of Patients With Aneurysmal Subarachnoid Hemorrhage. Stroke 55, 2305–2314 (2024) .

4. Li, Q., Lv, N., Li, L., Gu, Y., Xu, L., Lv, M., Huang, C., Mao, G., Lu, H., Zhong, S., et al. Intracranial Aneurysms Managed by Parent Artery Reconstruction Using Tubridge Flow Diverter study: 1-year outcomes. J Neurosurg 141, 1697–1704 (2024).

5. Marbacher, S., Grüter, B.E., Wanderer, S., Andereggen, L., Cattaneo, M., Trost, P., Gruber, P., Diepers, M., Remonda, L., Steiger, H.J. Risk of intracranial aneurysm recurrence after microsurgical clipping based on 3D digital subtraction angiography. J Neurosurg 29, 1–7 (2022) .

6. El Bakkouri, Y., Chidiac, R., Delisle, C., Corriveau, J., Cagnone, G., Gaonac’h-Lovejoy, V., Chin, A., Lécuyer, É., Angers, S., Joyal, J.S., et al. ZO-1 interacts with YB-1 in endothelial cells to regulate stress granule formation during angiogenesis. Nat Commun 15, 4405 (2024) .

7. Kataoka, K., Taneda, M., Asai, T., Kinoshita, A., Ito, M., Kuroda, R.Structural fragility and inflammatory response of ruptured cerebral aneurysms. a comparative study between ruptured and unruptured cerebral aneurysms. Stroke 30, 1396–1401 (1999) .

8. Ono, I., Abekura, Y., Kawashima, A., Oka, M., Okada, A., Hara, S., Miyamoto, S., Kataoka, H., Ishii, A., Yamamoto, K., et al.Endothelial cell malfunction in unruptured intracranial aneurysm lesions revealed using a 3D-casted mold. J Neuropathol Exp Neurol 82, 49–56 (2022) .

9. Liu, Q., Zhang, Y., Yang, J., Yang, Y., Li, M., Chen, S., Jiang, P., Wang, N., Zhang, Y., Liu, J., et al.The relationship of morphological-hemodynamic characteristics, inflammation, and remodeling of aneurysm wall in unruptured intracranial aneurysms. Transl Stroke Res 13,88–99 (2022) .

10. Hasan, D.M., Chalouhi, N., Jabbour, P., Magnotta, V.A., Kung, D.K., Young, W.L.Imaging aspirin effect on macrophages in the wall of human cerebral aneurysms using ferumoxytol-enhanced MRI: preliminary results. J Neuroradiol 40, 187–191 (2013) .

11. Bakker, M.K., van der Spek, R.A.A., van Rheenen, W., Morel, S., Bourcier, R., Hostettler, I.C., Alg V.S., van Eijk, K.R., Koido, M., Akiyama, M., et al. Genome-wide association study of intracranial aneurysms identifies 17 risk loci and genetic overlap with clinical risk factors. Nat Genet 52, 1303–1313 (2020).

12. Wu, C., Liu, H., Zuo, Q., Jiang, A., Wang, C., Lv, N., Lin, R., Wang, Y., Zong, K., Wei, Y., et al. Identifying novel risk genes in intracranial aneurysm by integrating human proteomes and genetics. Brain 147, 2817–2825(2024) .

13. Rinkel, G. J. E., Ruigrok, Y. M., Krings, T., Etminan, N., & Vergouwen, M. D. I. Advances in screening and management of unruptured intracranial aneurysms. The Lancet. Neurology, 24, 958–968 (2025).

14. Chen, H., McIntyre, M. K., Kakadiya, J., Rai, P., Essibayi, M. A., Salim, H. A., Azzam, A. Y., Latifikhereshky, S., Yedavalli, V. S., Altschul, D. J., et al. GLP-1 Receptor Agonist Use Is Associated With Lower Risk of Intracranial Aneurysm Rupture and Rupture Severity. Stroke, 57, 1979–1990 (2026).

15. Kolega, J., Gao, L., Mandelbaum, M., Mocco, J., Siddiqui, A.H., Natarajan, S.K., Meng, H. Cellular and molecular responses of the basilar terminus to hemodynamics during intracranial aneurysm initiation in a rabbit model. J Vasc Res 48,429–442 (2011).

16. Kaneko, N., Mashiko, T., Namba, K., Tateshima, S., Watanabe, E., Kawai, K. A patient-specific intracranial aneurysm model with endothelial lining: a novel *in vitro* approach to bridge the gap between biology and flow dynamics. J Neurointerv Surg 10, 306–309 (2018).

17. Aoki, T., Nishimura, M., Matsuoka, T., Yamamoto, K., Furuyashiki, T., Kataoka, H., Kitaoka, S., Ishibashi, R., Ishibazawa, A., Miyamoto, S., et al. PGE(2) -EP(2) signalling in endothelium is activated by haemodynamic stress and induces cerebral aneurysm through an amplifying loop via NF-κB. Br J Pharmacol 163,1237–1249 (2011).

18. Aoki, T., Yamamoto, K., Fukuda, M., Shimogonya, Y., Fukuda, S., Narumiya, S.Sustained expression of MCP-1 by low wall shear stress loading concomitant with turbulent flow on endothelial cells of intracranial aneurysm. Acta Neuropathol Commun 4, 48 (2016) .

19. Wang, Z., Kolega, J., Hoi, Y., Gao, L., Swartz, D.D., Levy, E.I., Mocco, J., Meng, H. Molecular alterations associated with aneurysmal remodeling are localized in the high hemodynamic stress region of a created carotid bifurcation. Neurosurgery 65, 169–178 (2009) .

20. Crouch, E.E., Bhaduri, A., Andrews, M.G., Cebrian-Silla, A., Diafos, L.N., Birrueta, J.O., Wedderburn-Pugh, K., Valenzuela, E.J., Bennett, N.K., Eze, U.C., et al. Ensembles of endothelial and mural cells promote angiogenesis in prenatal human brain. Cell 185, 3753–3769 (2022) .

21. Serafin, D.S., Harris, N.R., Bálint, L., Douglas, E.S., Caron, K.M. Proximity interactome of lymphatic VE-cadherin reveals mechanisms of junctional remodeling and reelin secretion. Nat Commun 15, 7734 (2024) .

22. Taylor, R.R., Keane, R.W., Guardiola, B., Martí, R., Alegre, D., Dietrich, W.D., Perez-Barcena, J., de Rivero Vaccari, J.P. Acute Neurovascular Inflammatory Profile in Patients with Aneurysmal Subarachnoid Hemorrhage. Biomolecules 15, 613 (2025).

23. Zhu, H., Zeng, Y., Tan, J., Li, M., Zhao, Y. HMGB1 Induced Oxidative Stress and Inflammation in Endothelial Cells Exposed to Impinging Flow. Cerebrovasc Dis 53, 437–448 (2024) .

24. Kamińska, J., Maciejczyk, M., Ćwiklińska, A., Matowicka-Karna, J., Koper-Lenkiewicz, O.M. Pro-Inflammatory and Anti-Inflammatory Cytokines Levels are Significantly Altered in Cerebrospinal Fluid of Unruptured Intracranial Aneurysm (UIA) Patients. J Inflamm Res 15, 6245–6261(2022) .

25. Kanematsu, Y., Kanematsu, M., Kurihara, C., Tada, Y., Tsou, T.L., van Rooijen, N., Lawton, M.T., Young, W.L., Liang, E.I., Nuki, Y., et al. Critical roles of macrophages in the formation of intracranial aneurysm. Stroke 42,173–178 (2011).

26. Hasan, D., Hashimoto, T., Kung, D., Macdonald, R.L., Winn, H.R., Heistad, D. Upregulation of cyclooxygenase-2 (COX-2) and microsomal prostaglandin E2 synthase-1 (mPGES-1) in wall of ruptured human cerebral aneurysms: preliminary results. Stroke 43, 1964–1967 (2012) .

27. Milanesi, E., Manda, G., Dobre, M., Codrici, E., Neagoe, I.V., Popescu, B.O., Bajenaru, O.A., Spiru, L., Tudose, C., Prada, G.I., et al.Distinctive Under-Expression Profile of Inflammatory and Redox Genes in the Blood of Elderly Patients with Cardiovascular Disease. J Inflamm Res 14, 429–442 (2021) .

28. Hennigs, J. K., Lüneburg, N., Stage, A., Schmitz, M., Körbelin, J., Harbaum, L., Matuszcak, C., Mienert, J., Bokemeyer, C., Böger, R. H., et al. The P2-receptor-mediated Ca2+ signalosome of the human pulmonary endothelium - implications for pulmonary arterial hypertension. Purinergic signalling, 15, 299–311 (2019).

29. Piazza, M., Guillemette, J., Dieckmann, T. Dynamics of nitric oxide synthasecalmodulin interactions at physiological calcium concentrations. Biochemistry 54, 1989–2000 (2015) .

30. Piazza, M., Dieckmann, T., Guillemette, J. Structural studies of a complex between endothelial nitric oxide synthase and calmodulin at physiological calcium concentration. Biochemistry 55,5962–5971 (2016) .

31. Liu, T.T., Xu, H.H., Liu, Z.J., Zhang, H.P., Zhou, H.T., Zhu, Z.X., Wang, Z.Q., Xue, J.Y., Li, Q., Ma, Y., et al. Downregulated calmodulin expression contributes to endothelial cell impairment in diabetes. Acta Pharmacol Sin 44, 2492–2503 (2023) .

