## supplementary figures for "Pharmacological up-regulation of CALM1 restore endothelial cells in intracranial aneurysm pathogenesis"

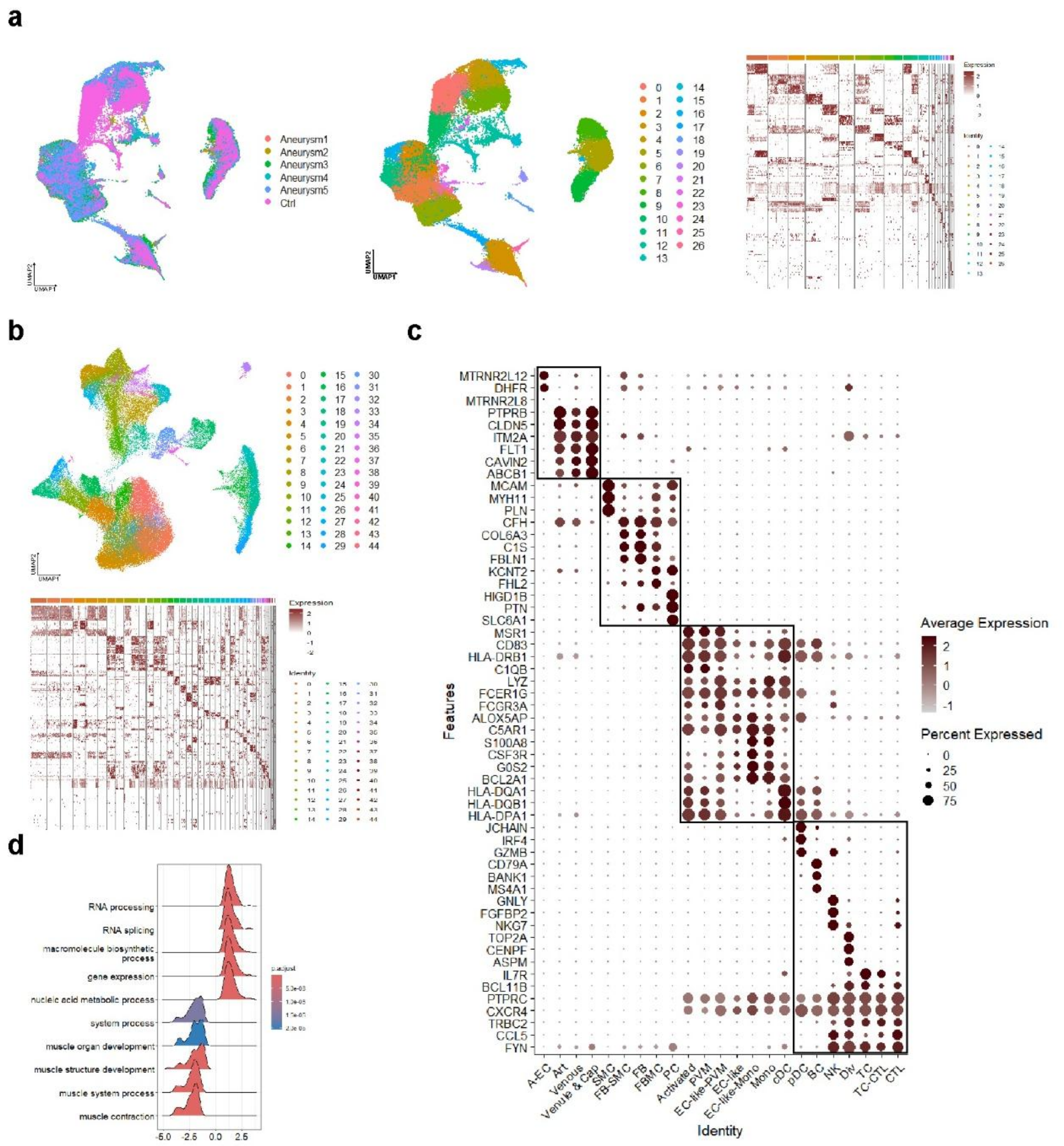


**Figure S1. Multi-omics analysis reveals cellular senescence as a key factor in IA development.**

a, UMAP visualization of clustering of cells from human IA samples and Ctrl sample after Seurat integration (left and middle). Heatmap showing marker genes of each cluster of cells.

b, UMAP visualization of clustering of cells from human IA samples, Ctrl sample, and cells from adult human cerebrovasculature scRNA-seq reference dataset after Seurat integration (top). Heatmap showing marker genes of each cluster of cells (bottom).

c, Dotplot showing markers of each cell type in cells from human IA samples, Ctrl sample, and cells from adult human cerebrovasculature scRNA-seq reference dataset after Seurat integration.

d, Ridgeplot showing Gene Ontology enrichment analysis of DEPs between human IA samples and Ctrl samples from proteomic result.

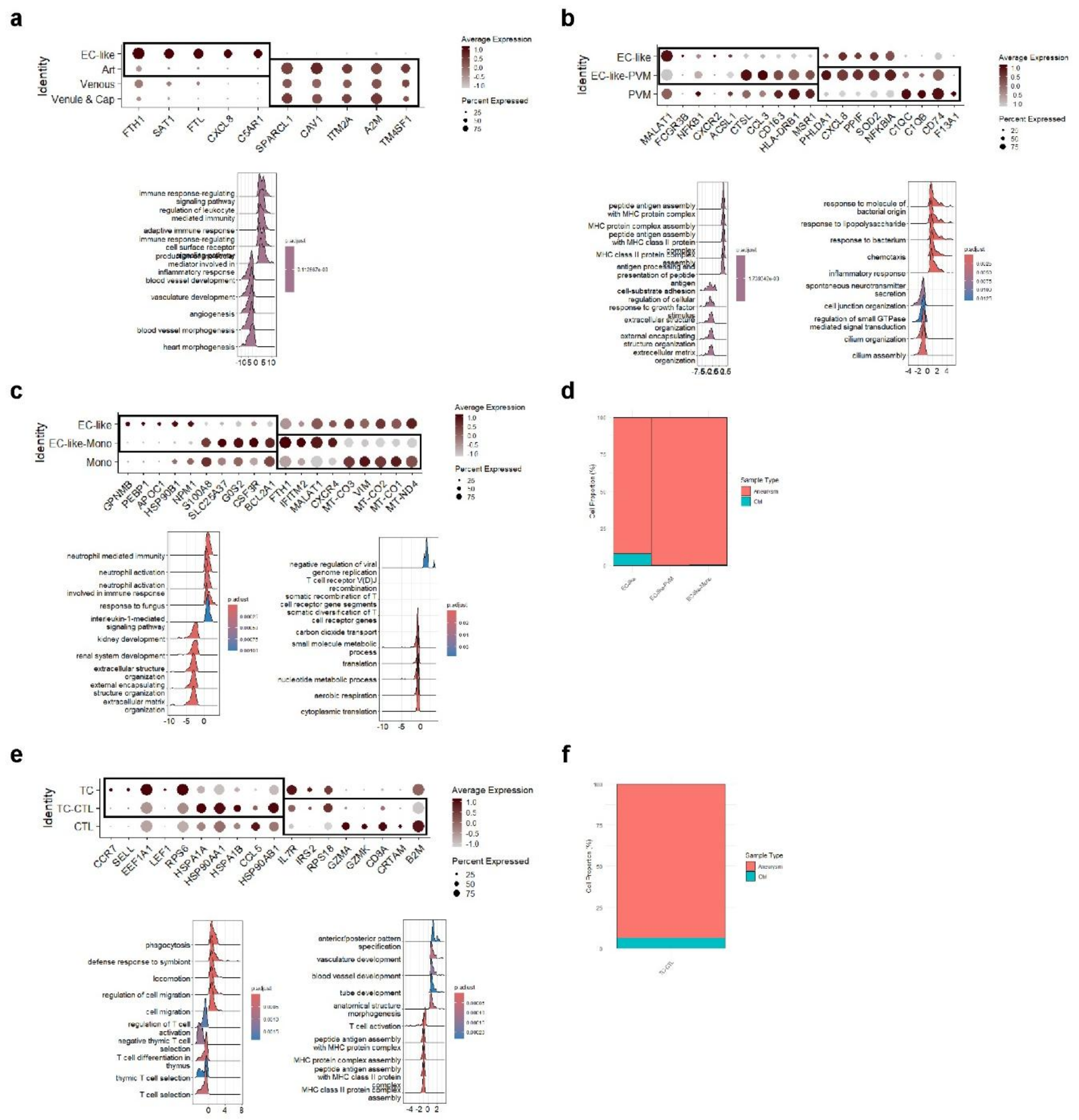

**Figure S2. Immune cells display cell fate shift, impaired cellular functions, and dysregulated immune responses in IAs.**

a, Dotplot showing DEGs between EC-like cells and other endothelial cells (top). Ridge plot showing Gene Ontology enrichment analysis of DEGs between EC-like cells and other endothelial cells (bottom).

b, Dotplot showing DEGs between EC-like-PVM cells and EC-like cells and PVM cells (top). Ridge plot showing Gene Ontology enrichment analysis of DEGs between EC-like-PVM cells and EC-like cells (bottom left). Ridge plot showing Gene Ontology enrichment analysis of DEGs between EC-like-PVM cells and PVM cells (bottom right).

c, Dotplot showing DEGs between EC-like-Mono cells and EC-like cells and Mono cells (top). Ridge plot showing Gene Ontology enrichment analysis of DEGs between EC-like-Mono cells and EC-like cells (bottom left). Ridge plot showing Gene Ontology enrichment analysis of DEGs between EC-like-Mono cells and Mono cells (bottom right).

d, Quantification of distribution of EC-like cells, EC-like-PVM cells and EC-like-Mono cells in human IA samples and Ctrl sample.

e, Dotplot showing DEGs between TC-CTL cells and T cells and CTL cells (top). Ridge plot showing Gene Ontology enrichment analysis of DEGs between TC-CTL cells cells and T cells (bottom left). Ridge plot showing Gene Ontology enrichment analysis of DEGs between TC-CTL cells and CTL cells (bottom right).

f, Quantification of distribution of TC-CTL cells in human IA samples and Ctrl sample.

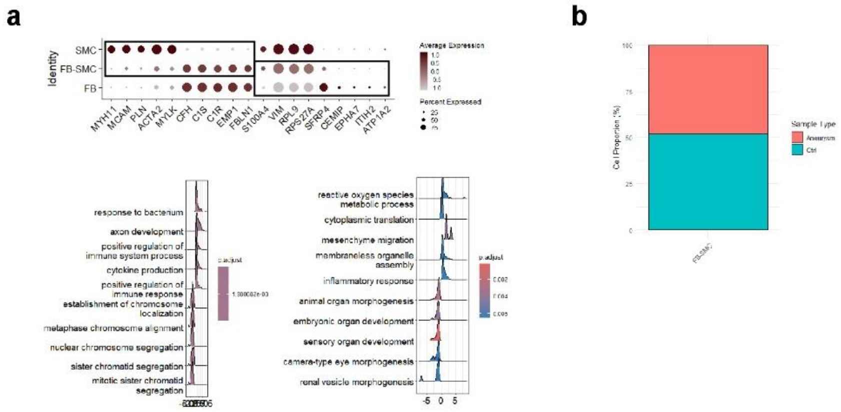


**Figure S3. Perivascular cells undergo pathological changes that disrupt vascular homeostasis in IAs.**

a, Dotplot showing DEGs between FB-SMC cells and SMC cells and FB cells (top). Ridge plot showing Gene Ontology enrichment analysis of DEGs between FB-SMC cells and SMC cells (bottom left). Ridge plot showing Gene Ontology enrichment analysis of DEGs between FB-SMC cells and FB cells (bottom right).

b, Quantification of distribution of FB-SMC cells in human IA samples and Ctrl sample.
